# ATF6 enables pathogen infection in ticks by inducing *stomatin* and altering cholesterol dynamics

**DOI:** 10.1101/2025.01.08.632023

**Authors:** Kaylee A. Vosbigian, Sarah J. Wright, Kristin L. Rosche, Elis A. Fisk, Elisabeth Ramirez-Zepp, Eric A. Shelden, Dana K. Shaw

## Abstract

How tick-borne pathogens interact with their hosts has been primarily studied in vertebrates where disease is observed. Comparatively less is known about pathogen interactions within the tick. Here, we report that *Ixodes scapularis* ticks infected with either *Anaplasma phagocytophilum* (causative agent of anaplasmosis) or *Borrelia burgdorferi* (causative agent of Lyme disease) show activation of the ATF6 branch of the unfolded protein response (UPR). Disabling ATF6 functionally restricts pathogen survival in ticks. When stimulated, ATF6 functions as a transcription factor, but is the least understood out of the three UPR pathways. To interrogate the *Ixodes* ATF6 transcriptional network, we developed a custom R script to query tick promoter sequences. This revealed *stomatin* as a potential gene target, which has roles in lipid homeostasis and vesical transport. *Ixodes stomatin* was experimentally validated as a bona fide ATF6-regulated gene through luciferase reporter assays, pharmacological activators, and RNAi transcriptional repression. Silencing *stomatin* decreased *A. phagocytophilum* colonization in *Ixodes* and disrupted cholesterol dynamics in tick cells. Furthermore, blocking *stomatin* restricted cholesterol availability to the bacterium, thereby inhibiting growth and survival. Taken together, we have identified the *Ixodes* ATF6 pathway as a novel contributor to vector competence through Stomatin-regulated cholesterol homeostasis. Moreover, our custom, web-based transcription factor binding site search tool “ArthroQuest” revealed that the ATF6-regulated nature of *stomatin* is unique to blood-feeding arthropods. Collectively, these findings highlight the importance of studying fundamental processes in non-model organisms.

**IMPORTANCE:** Host-pathogen interactions for tick-borne pathogens like *Anaplasma phagocytophilum* (causative agent of Anaplasmosis) have been primarily studied in mammalian hosts. Comparatively less is known about interactions within the tick. Herein, we find that tick-borne pathogens activate the cellular stress response receptor, ATF6, in *Ixodes* ticks. Upon activation, ATF6 is cleaved and the cytosolic portion translocates to the nucleus to function as a transcription factor that coordinates gene expression networks. Using a custom script in R to query the *Ixodes* ATF6 regulome, *stomatin* was identified as an ATF6-regulated target that supports *Anaplasma* colonization by facilitating cholesterol availability to the bacterium. Moreover, our custom, web-based tool “ArthroQuest” found that the ATF6-regulated nature of *stomatin* is unique to arthropods. Given that lipid hijacking is common among arthropod-borne microbes, ATF6-mediated induction of *stomatin* may be a mechanism that is exploited in many vector-pathogen relationships for the survival and persistence of transmissible microbes. Collectively, this study identified a novel contributor to vector competence and highlights the importance of studying molecular networks in non-model organisms.

## INTRODUCTION

The North American deer tick, *Ixodes scapularis*, can transmit up to seven different pathogens that impact human and animal health including *Anaplasma phagocytophilum* (causative agent of Anaplasmosis) and *Borrelia burgdorferi* (causative agent of Lyme Disease)^1^. The continuous rise in reported cases of tick-borne disease^2–10^ underscores the need for novel intervention strategies. Although the intricacies of mammalian host-pathogen interactions have been well-studied, comparatively little is known about tick-pathogen interactions.

Recently we have shown that *A. phagocytophilum* and *B. burgdorferi* activate the unfolded protein response (UPR) in ticks, which influences microbial colonization and persistence in the arthropod^11,12^. The UPR is a cellular response network that is initiated by three endoplasmic reticulum (ER) transmembrane receptors IRE1α, PERK, and ATF6. Each branch of the UPR initiates a signaling cascade and coordinates gene expression networks by activating specific transcription factors. We have shown that the IRE1α-TRAF2 pathway leads to microbe-restricting immune responses in arthropods by activating the NF-κB-like molecule, Relish^11^. We have also demonstrated that stimulating PERK activates the antioxidant transcription factor, Nrf2, which facilitates pathogen persistence in ticks^12^. Out of the three UPR receptors, ATF6 is the least understood^13^. When activated, site-1 and site-2 proteases cleave the cytosolic portion of ATF6, which allows it to translocate to the nucleus and act as a transcriptional regulator (nATF6)^14^. The role of ATF6 has never been explored in arthropod vectors.

Here, we demonstrate that *Ixodes* ATF6 is activated by tick-borne pathogens and supports *A. phagocytophilum* colonization in ticks. To determine how ATF6 impacts vector competence, we used protein modeling and a custom transcription factor binding site query to probe the ATF6 regulatory network in *I. scapularis.* Gene ontology (GO) and Reactome analyses identified Stomatin, a lipid homeostasis and vesical transport protein, as a potential gene regulated by ATF6 in ticks. Using pharmacological manipulations, RNA interference (RNAi), and quantitative fluorescent assays, we found that Stomatin supports pathogen colonization in ticks by facilitating cholesterol acquisition by the bacterium. These findings demonstrate that *stomatin* is induced during the arthropod-phase of the pathogen life cycle to enable survival and persistence in the vector.

Programs that predict transcription factor regulatory networks are generally restricted to model organisms, leaving out many arthropod vectors. We used our custom R script to develop a publicly available, web-based tool termed “ArthroQuest” that currently allows users to query 20 different arthropod vector genomes, in addition to *Drosophila* and humans. Queries with ArthroQuest revealed that the ATF6-regulated nature of *stomatin* appears to be unique to arthropods. Given that lipid hijacking and cholesterol incorporation is common in many arthropod-borne microbes^15^, ATF6-mediated induction of *stomatin* may be a shared phenomenon among many vector-pathogen relationships that is exploited for the survival and persistence of transmissible pathogens.

## RESULTS

### ATF6 is induced by tick-borne pathogens and supports microbial survival

Previously, we found that *A. phagocytophilum* and *B. burgdorferi* activate the UPR through ER receptors IRE1α and PERK, which influence bacterial colonization of the tick^11,12^. The UPR coordinates a variety of transcription factors that modulate the expression of genes involved in cellular stress responses, immunity, and several other physiological conditions^16–18^. To determine which UPR-associated transcription factors respond to infection, we employed a surrogate luciferase reporter plasmid assay^12^ and found that ATF6 was significantly activated by both *A. phagocytophilum* and *B. burgdorferi* (Fig 1A-B). We next asked if *atf6* is induced in ticks during infection. *I. scapularis* larvae were fed to repletion on either *A. phagocytophilum* or *B. burgdorferi-* infected mice and gene expression was analyzed by quantitative real-time PCR (qRT-PCR). In both groups of infected larvae, *atf6* expression was significantly increased (Fig 1C-D). To determine how this may impact pathogen colonization, we silenced *atf6* in tick cells using RNAi prior to infecting with *A. phagocytophilum* and found that knocking down *atf6* reduced bacterial survival and replication in tick cells (Fig 1E-F). Collectively, this indicates that ATF6 is activated during infection and supports pathogen survival.

**Figure 1.**
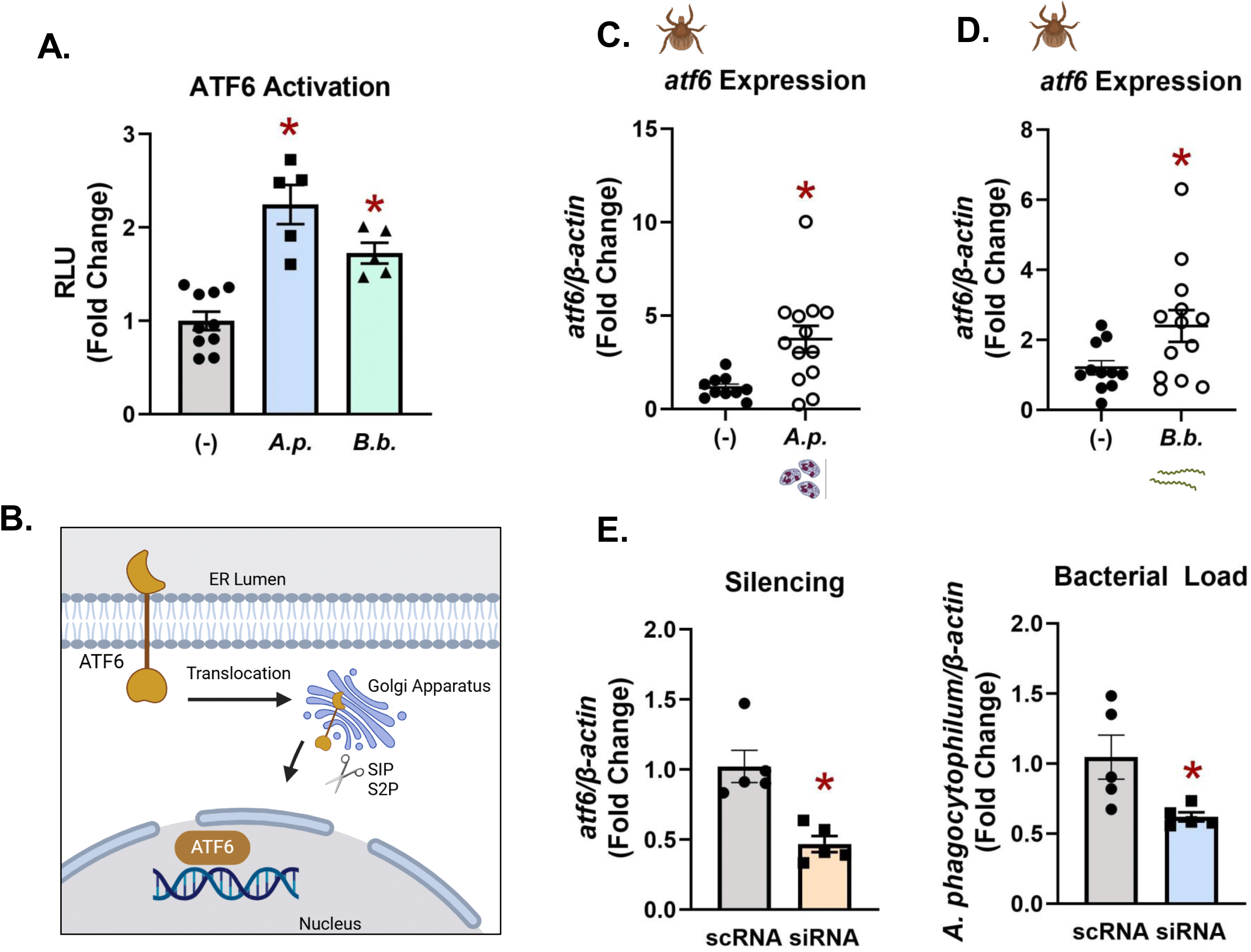
ATF6 is induced by tick-borne pathogens during infection and supports vector colonization. (A) HEK 293T cells were transfected with an ATF6 luciferase reporter plasmid. Transfected cells were infected with *A. phagocytophilum* (MOI 50) or *B. burgdorferi* (MOI 200) for 18 hours and normalized to no infection (-). RLU is relative luminescence units. (B) Schematic depicting ATF6 activation. (C-D) Gene expression of *atf6* in *I. scapularis* larvae rested for 7 days after feeding on *Anaplasma-*infected mice (C) and rested for 14 days after feeding on *Borrelia-*infected mice (D). Expression was assessed via qRT-PCR. Each data point is representative of 1 larva. (-) is no infection. *A.p.* is *A. phagocytophilum*. *B.b.* is *B. burgerdorferi.* (E) RNAi knockdown of *atf6* in ISE6 cells for 5 days and infected with *A. phagocytophilum* (MOI 50). scRNA, scrambled RNA; siRNA, small interfering RNA. Experiments are representative of at least 2 experimental replicates. *P < 0.05.

### Identifying genes putatively regulated by ATF6 in ticks

Out of the three UPR pathways, ATF6 and the gene network it coordinates is the least understood^13^. To examine how ATF6 is mechanistically impacting pathogen dynamics in the tick, we first aligned sequences from *I. scapularis* and humans. We found that, although there was low overall conservation (31.56% identity), there was good conservation in the basic leucine zipper (bZIP) domain which is the DNA-interacting portion of ATF6 (Supplementary Fig 1A; Supplementary Table 1). Moreover, the amino acid residues defined as “essential for DNA binding”, K304, N305, and R306, were 100% conserved between the two sequences^19^. We next used AlphaFold to predict the protein structure of *Ixodes* ATF6 and then aligned it with the human structure^20,21^ using ChimeraX^22^ (Fig 2A, Supplementary Fig 1B). We found that the bZIP domain was structurally well-conserved and that the DNA-binding amino acid residues were found in the same orientation between the two proteins (Fig 2B).

**Figure 2.**
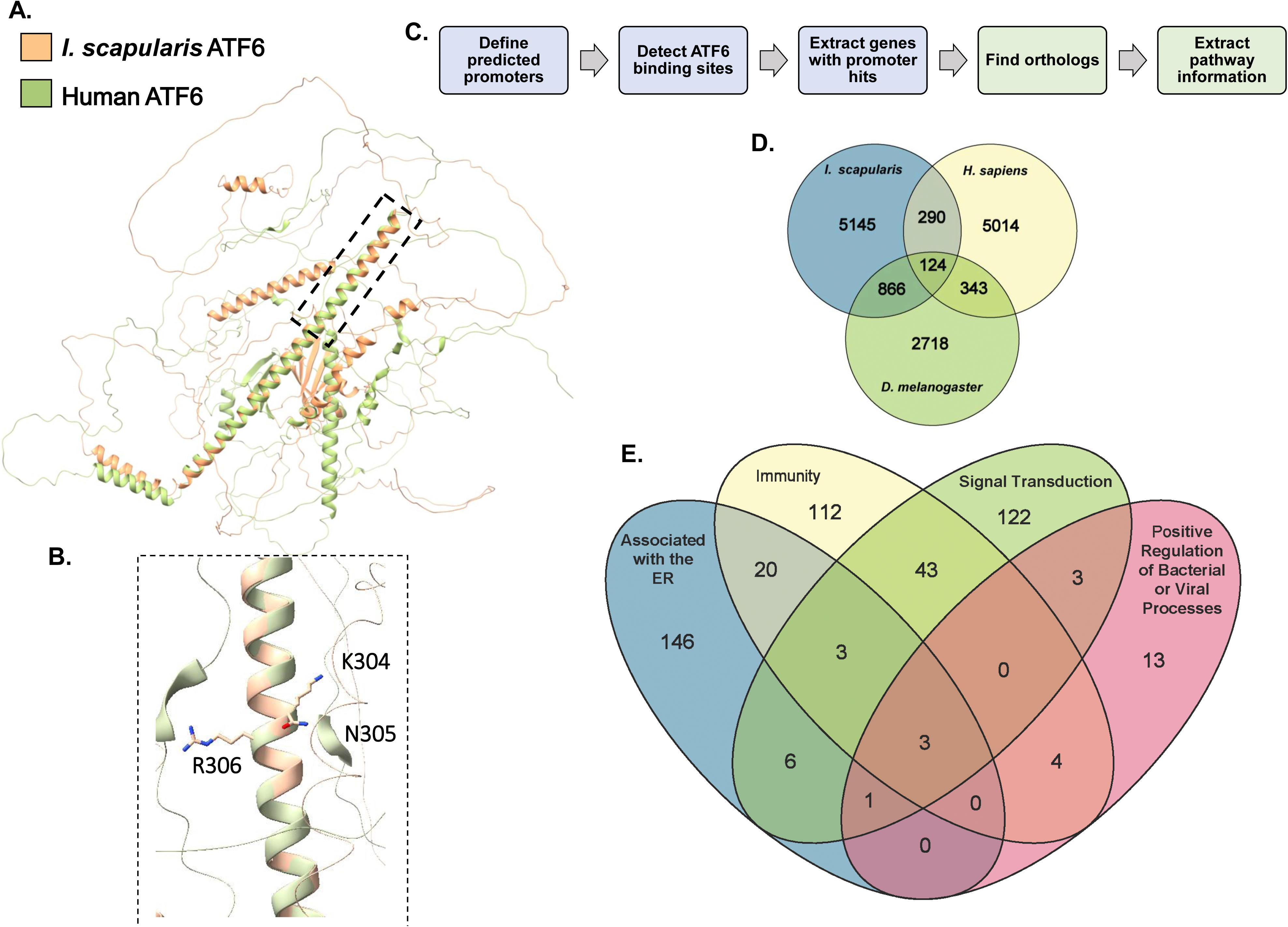
Predicting the ATF6 regulatory network in ticks. (A) The *I. scapularis* ATF6 (orange) predicted with AlphaFold has a conserved DNA binding domain compared to the human ortholog (green). (B) A portion of the bZIP domain and the DNA binding residues on ATF6 are visualized. (C) Schematic of the developed R script. (D) Comparison of genes predicted to be regulated by ATF6 from the *I. scapularis*, *D. melanogaster,* and *H. sapiens*. (E) GO terms and Reactome pathways represented in predicted ATF6-regulated genes from *I. scapularis* genome.

The conservation in DNA-binding domain structure and residues between humans and ticks provided the impetus to interrogate the *Ixodes* genome for ATF6-regulated genes. Non-model organisms, such as ticks, have a limited number of genome and proteome analysis tools. Moreover, the tools that are available are often not well-developed. To circumvent this issue, we created a custom query in R to search for ATF6-regulated genes in *I. scapularis* (Fig 2C). Predicted promoter regions in the *I. scapularis* genome were defined as 1,000 base pairs upstream from all coding regions. We then searched for ATF6-binding motifs in all promoters. ATF6 functions as a homodimer that promotes gene expression either by itself or in combination with the general transcription factor NF-Y (nuclear transcription factor Y). By itself, ATF6 can drive gene expression by binding TGACGTG within a promoter sequence^19^. In combination, ATF6 and NF-Y can drive transcription by binding CCACG and CCAAT, respectively^23,24^. We scanned all predicted *Ixodes* promoters for either TGACGTG or CCACG in the presence of CCAAT. For comparison, we also predicted ATF6-binding sites in humans and the model insect *D. melanogaster.* Genes downstream from promoters containing ATF6-binding sites were compiled, identified, and compared (Fig 2D; Supplementary Table 2). This revealed that known ATF6-regulated genes, including *binding immunoglobulin protein* (*BiP*) and *x-box binding protein 1* (*xbp1*), were present in all three organisms, demonstrating that our custom R script correctly identifies ATF6-regulated genes. We also observed that more genes were shared between *Ixodes* and *Drosophila* (866) than humans and *Ixodes* (290), but the large majority of genes were unique to each organism (Fig 2D).

We next sought to analyze the ATF6-regulated gene network in ticks. However, when compared to humans or *Drosophila*, relatively little gene information is available for *Ixodes.* To overcome this obstacle, we identified human and *Drosophila* orthologs for tick gene targets and then extracted corresponding gene information (Supplementary Table 2). 71% of the *Ixodes* genes putatively regulated ATF6 mapped to human and/or *Drosophila* orthologs. Corresponding Gene Ontology (GO) and Reactome information^25–27^ were used to perform pathway enrichment analysis^28^ (Supplementary Fig 2), which revealed that GO terms associated with the UPR were enriched (Supplementary Table 3). This also returned categories of interest with the potential to impact microbial colonization including “Immunity”, “Positive Regulation of Bacterial or Viral Processes”, “Signal Transduction”, and “Associated with the ER”^14,29^. From *Ixodes*, we found 185 genes associated with Immunity (GO:0006955, R-HSA-768256), 24 with Positive Regulation of Bacterial or Viral Processes (GO:0048524, GO:1900425), 181 with Signal Transduction (R-HSA-162582), and 179 Associated with the ER (GO:0005783) (Fig 2E). The genes *stomatin, neurogenic locus notch homolog protein 1 (notch1),* and *protein disulfide isomerase (pdi)* were found in all four GO categories. *Pdi* is a known ATF6-regulated gene in mammals and assists in the formation of disulfide bonds^30^. *Notch1* is also known to be regulated by ATF6 and is involved in immunity, cellular development, and apoptosis^31,32^. Stomatin has roles in lipid homeostasis and vesical transport^33^, but has never been linked to ATF6. Since *A. phagocytophilum* incorporates lipids and cholesterol into its membrane, we hypothesized that regulation of *stomatin* expression could be how ATF6 is supporting *A. phagocytophilum* in ticks.

### Tick stomatin is upregulated by ATF6 during infection

We next used AlphaFold3^34^ to predict structural interactions between ATF6 and the *stomatin* promoter (Fig 3A; Supplementary Fig 3A). We found that the *Ixodes* ATF6 homodimer (Fig 3A, tan) and DNA-interacting residues K304, N305, and R306 (Fig. 3A, purple) were predicted to be in direct contact with the ATF6-binding nucleotide motif found in the *stomatin* promoter, CCACG (Fig 3A, yellow). To test whether ATF6 activation increases *stomatin,* we used a drug, AA147, that selectively activates ATF6 independent of ER stress^35^. AA147 did not impact cell viability (Supplementary Fig 3B), but did cause a dose-dependent increase in *stomatin* expression (Fig 3B). Since pharmacological manipulators can have off-target effects, we also silenced *atf6* in tick cells using RNAi. We found that decreasing *atf6* levels (Fig 1E) caused a significant decline in *stomatin* expression (Fig 3C), altogether demonstrating that ATF6 positively correlates with *stomatin* expression.

**Figure 3.**
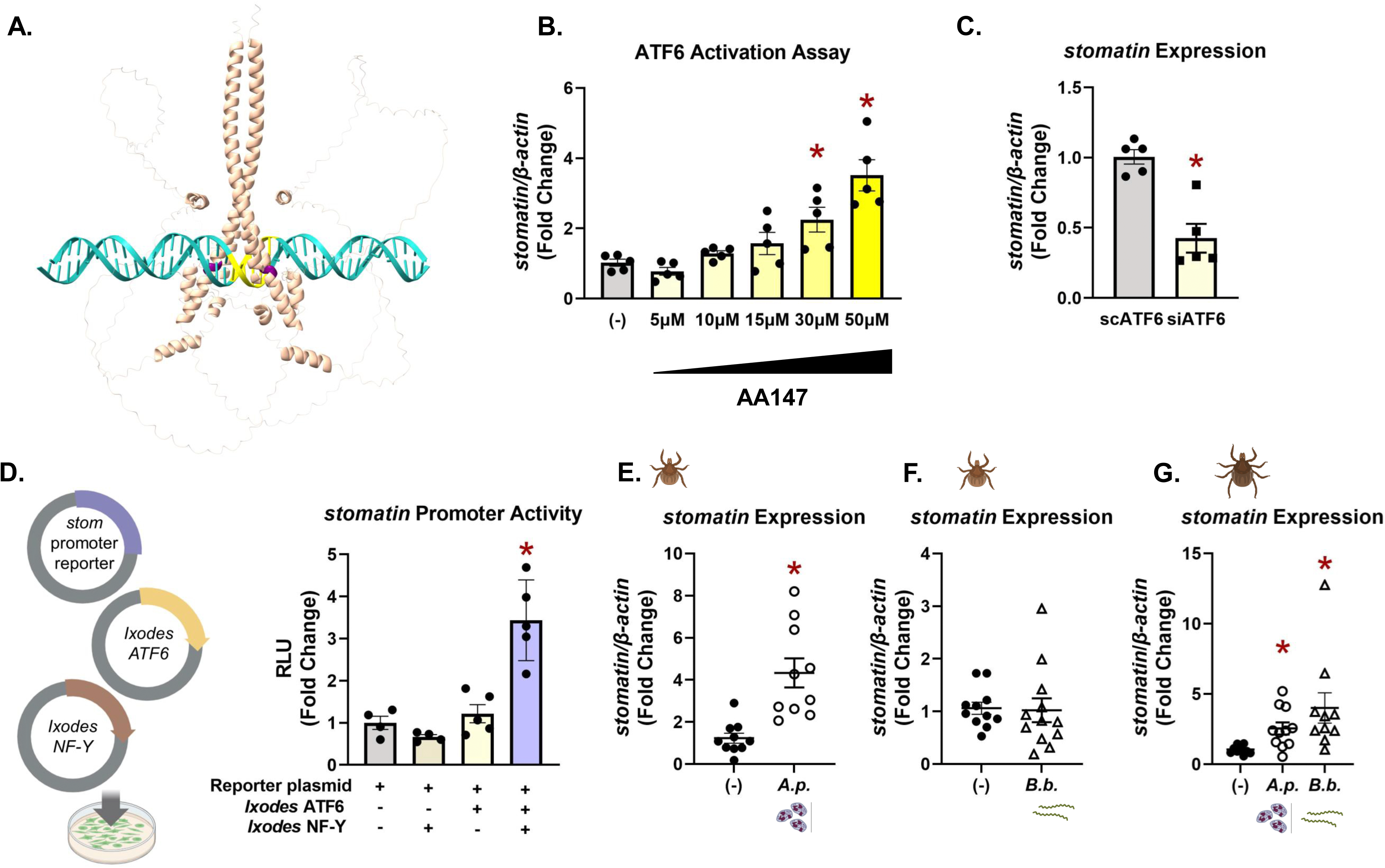
ATF6 upregulates *stomatin* during infection in ticks. (A) The AlphaFold3 predicted DNA-protein complex with *I. scapularis* nATF6 (tan) and the predicted promoter of *stomatin* (teal). The ATF6 binding site is highlighted in yellow and DNA binding residues highlighted in purple. (B-C) *stomatin* expression quantified from (B) ISE6 cells treated with 5 µM-50 µM of AA147 for 24 hours and (C) ISE6 cells transfected with siRNA targeting *atf6* or a scrambled control. (D) HEK 293T cells were co-transfected with plasmids constitutively expressing *I. scapularis* nATF6, *I. scapularis* NF-Y, and a luciferase reporter plasmid with the *stomatin* promoter. Luciferase activity was normalized to the control containing only the luciferase reporter plasmid. RLU is relative luminescence units. (E-F) Gene expression of larvae rested for 7 days after being fed on *Anaplasma-*infected mice (E) or rested for 14 days after being fed on *B. burgdorferi*-infected mice (F). (G) *stomatin* gene expression from unfed, infected *I. scapularis* nymphs. *A.p.* is *A. phagocytophilum*. *B.b.* is *B. burgerdorferi.* Expression assessed via qRT-PCR. *P < 0.05.

To experimentally validate that ATF6 binds the *stomatin* promoter and drives expression, we designed a Luciferase reporter assay. First, we cloned the *Ixodes stomatin* promoter upstream from a *luciferase* gene. Next, plasmids were constructed that constitutively expressed recombinant versions of *Ixodes* nATF6 and NF-Y (Supplementary Fig 4A-4B). All three plasmids were then co-transfected into HEK 293T cells for 24 hours (Fig 3D). A positive control plasmid containing *luciferase* driven by an *atf6* promoter was also co-transfected with *Ixodes* nATF6 and NF-Y constructs (Supplementary Fig 4C). After 24 hours, D-luciferin was added to the cells and Luciferase activity was assayed. We found a significant increase in Luciferase activity when cells were expressing both *Ixodes* nATF6 and NF-Y, indicating *stomatin* promoter activity (Fig 3D). Altogether, these data demonstrate that ATF6, in combination with the ubiquitously expressed transcription factor NF-Y, positively regulates the expression of *stomatin* in *Ixodes* ticks.

We next evaluated *in vivo stomatin* expression in infected ticks. *Stomatin* was quantified in larvae that had been fed on either *A. phagocytophilum* (Fig 3E) or *B. burgdorferi* (Fig 3F) infected mice, and then rested for 7 days or 14 days post-repletion, respectively. This time period correlates with the expansion of microbes in the tick post-repletion^36^. We also quantified *stomatin* expression in *A. phagocytophilum* or *B. burgdorferi* infected, flat nymphs (Fig 3G). From both life stages, we found that ticks infected with *A. phagocytophilum* had elevated levels of *stomatin* expression when compared to uninfected ticks. Ticks infected with *B. burgdorferi* had no differences at the rested larval stage, but significantly increased levels of *stomatin* at the unfed nymph stage. It is not clear why *stomatin* is induced in *A. phagocytophilum-*infected, rested larvae, but not *B. burgdorferi.* It is possible that differences in niche colonization and subcellular location between the two pathogens are responsible for this observation. This data, together with infection-induced ATF6 transcriptional activity (Fig 1D) demonstrates that ATF6 positively regulates *stomatin* during *A. phagocytophilum* and *B. burgdorferi* infection in ticks.

### Arthropod infection is supported by stomatin

Since Stomatin expression increased with *A. phagocytophilum* acquisition in ticks, we next asked what impact it has on microbial colonization. To address this, we used RNAi to silence *stomatin* in tick cells prior to infecting with *A. phagocytophilum*. We found that blocking *stomatin* expression caused a decrease in *Anaplasma* numbers (Fig 4A), similar to what was observed when silencing *atf6* (Fig 1E). To determine how Stomatin impacts microbial colonization *in vivo*, we immersed *Ixodes* larvae^11,12^ in siRNA targeting *stomatin.* Ticks were then allowed to feed to repletion on *A. phagocytophilum-*infected mice. We found a 2-fold decrease in *Anaplasma* numbers when *stomatin* expression was knocked down relative to the scrambled control (Fig 4B), demonstrating that *A. phagocytophilum* infection and colonization of ticks is supported by Stomatin.

**Figure 4.**
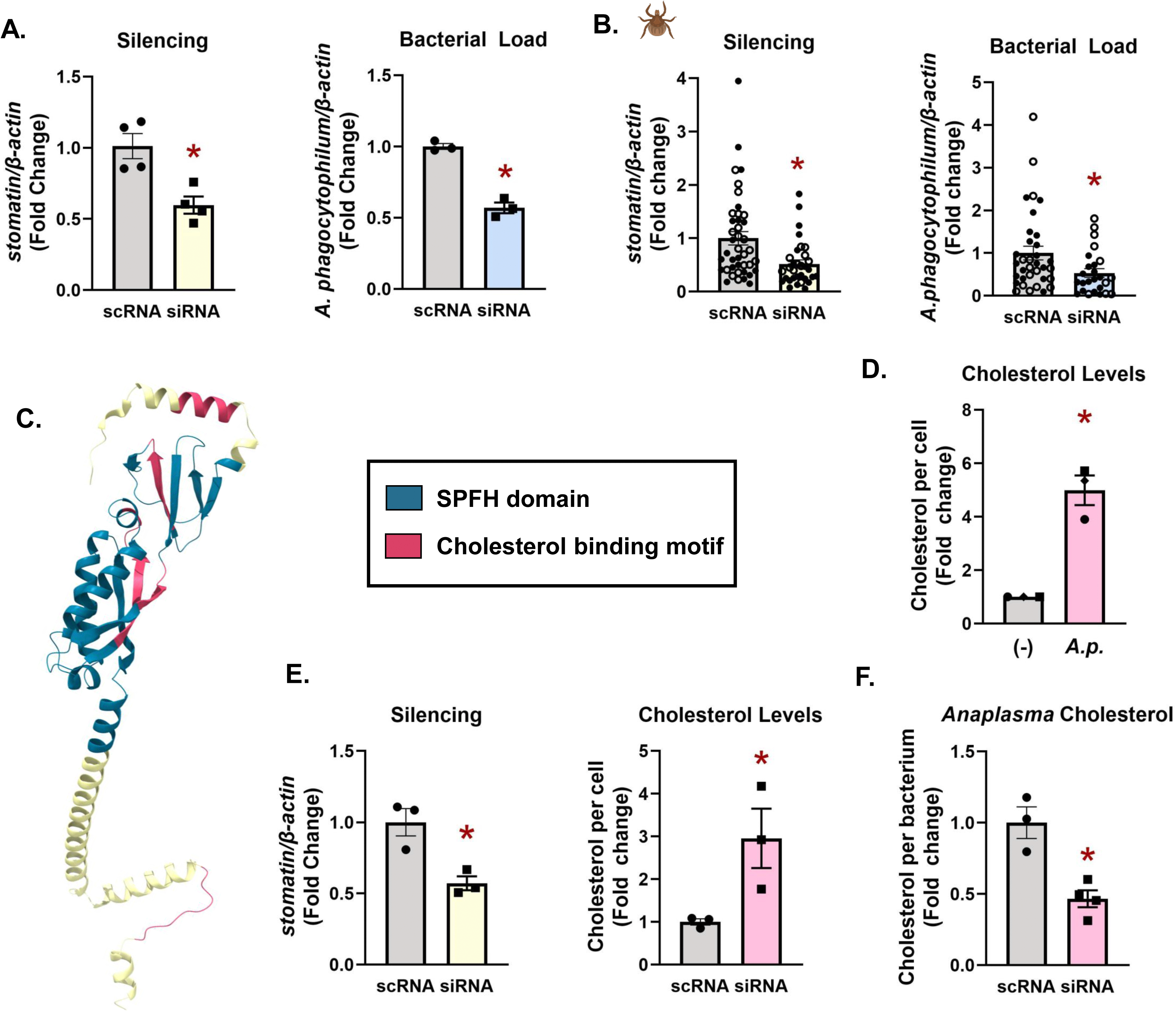
Stomatin supports infection by facillitating *A. phagocytophilum* cholesterol uptake. (A) ISE6 cells were transfected with siRNA or a scrambled control. Cells were infected with *A. phagocytophilum* (MOI 50) for 18 hours. (B) siRNA-treated or scrambled control-treated larvae were fed on *A. phagocytophilum-*infected mice. Each data point is representative of 1 larva. Open and closed dots represent experimental replicates 1 and 2. Gene silencing and bacterial burden were measured by qRT-PCR. (C) AlphaFold-predicted *I. scapularis* Stomatin with SPFH domain highlighted in blue and cholesterol binding domains highlighted in pink. (D) Total cholesterol quantified from *A. phagocytophilum-*infected ISE6 compared to uninfected cells (-). Each dot represents an experimental replicate. (E) Cholesterol quantified from cells treated with silencing RNAs targeting *stomatin* or scrambled controls. (F) Cholesterol quantified in *A. phagocytophilum* grown in *stomatin-*depleted ISE6 tick cells. Gene silencing was measured by qRT-PCR. scRNA, scrambled RNA; siRNA, small interfering RNA. Experiments are representative of at least 2 experimental replicates. *P < 0.05.

### Stomatin supports cholesterol incorporation into Anaplasma

Stomatin is a member of the stomatin/prohibitin/flotillin/HflK/C (SPFH) family found in lipid rafts on cell membranes, lipid droplets, and endosomes and is known to be involved in vesicle trafficking^37^. When comparing human and *Ixodes* Stomatin, we found that the amino acid sequences and structures are well conserved (Supplementary Fig 5A-C). The *Ixodes* Stomatin SPFH domain (residues 32-191) contains four cholesterol recognition/interaction amino acid consensus sequences, CRAC and CARC (Fig 4C, pink; Supplementary Fig 5D)^38^. Outside of the SPFH domain, there are two additional CARC domains. This suggests that tick Stomatin may bind cholesterol and function similarly to other SPFH family members^37^.

Cholesterol is essential to the development of *A. phagocytophilum* infection^39^. In mammals, *A. phagocytophilum* will intercept free cholesterol from the low-density lipoprotein uptake pathway^39^ and incorporate it for structural support of its membrane. We therefore asked if *A. phagocytophilum* impacts cholesterol dynamics in ticks. Using an Amplex Red cholesterol assay, cholesterol was quantified in uninfected tick cells relative to tick cells that were persistently infected with *A. phagocytophilum*. We found that infection was associated with significantly more cholesterol compared to uninfected cells, indicating that *Anaplasma* induces cholesterol accumulation in ticks (Fig 4D).

SPFH-domain containing proteins regulate cholesterol uptake, distribution within the cell, and exportation when levels become too high^40,37^. To understand how Stomatin may be impacting cholesterol dynamics within tick cells, we transfected silencing RNAs targeting *stomatin* into uninfected cells and then quantified cholesterol. We found that when *stomatin* expression is reduced there is an increase in total cholesterol, implying that Stomatin is involved in proper distribution, localization, and export of cholesterol from the cell (Fig 4E).

Given the *Anaplasma-*supporting role that Stomatin plays, we asked if it could be facilitating infection and colonization by shuttling cholesterol to the bacterium. To address this possibility, we silenced *stomatin* in persistently infected tick cells for five days (Supplementary Fig 5E). *A. phagocytophilum* were then mechanically lysed from tick cells, washed, and separated from host cell debris. Bacterial cholesterol levels were then quantified using the Amplex Red cholesterol assay. We found that when *A. phagocytophilum* is grown in cells with depleted Stomatin, there is significantly less total cholesterol incorporated into the bacteria compared to non-silenced controls (Fig 4F). Therefore, although knocking down *stomatin* caused increased total cellular cholesterol in tick cells, our data shows that it is not accessible by *Anaplasma.* This finding depicts a model where *Anaplasma* activates ATF6 in *Ixodes*, which upregulates Stomatin expression and functionally supports the microbe by facilitating cholesterol delivery, likely for cell wall structure and growth (Fig 5A).

**Figure 5.**
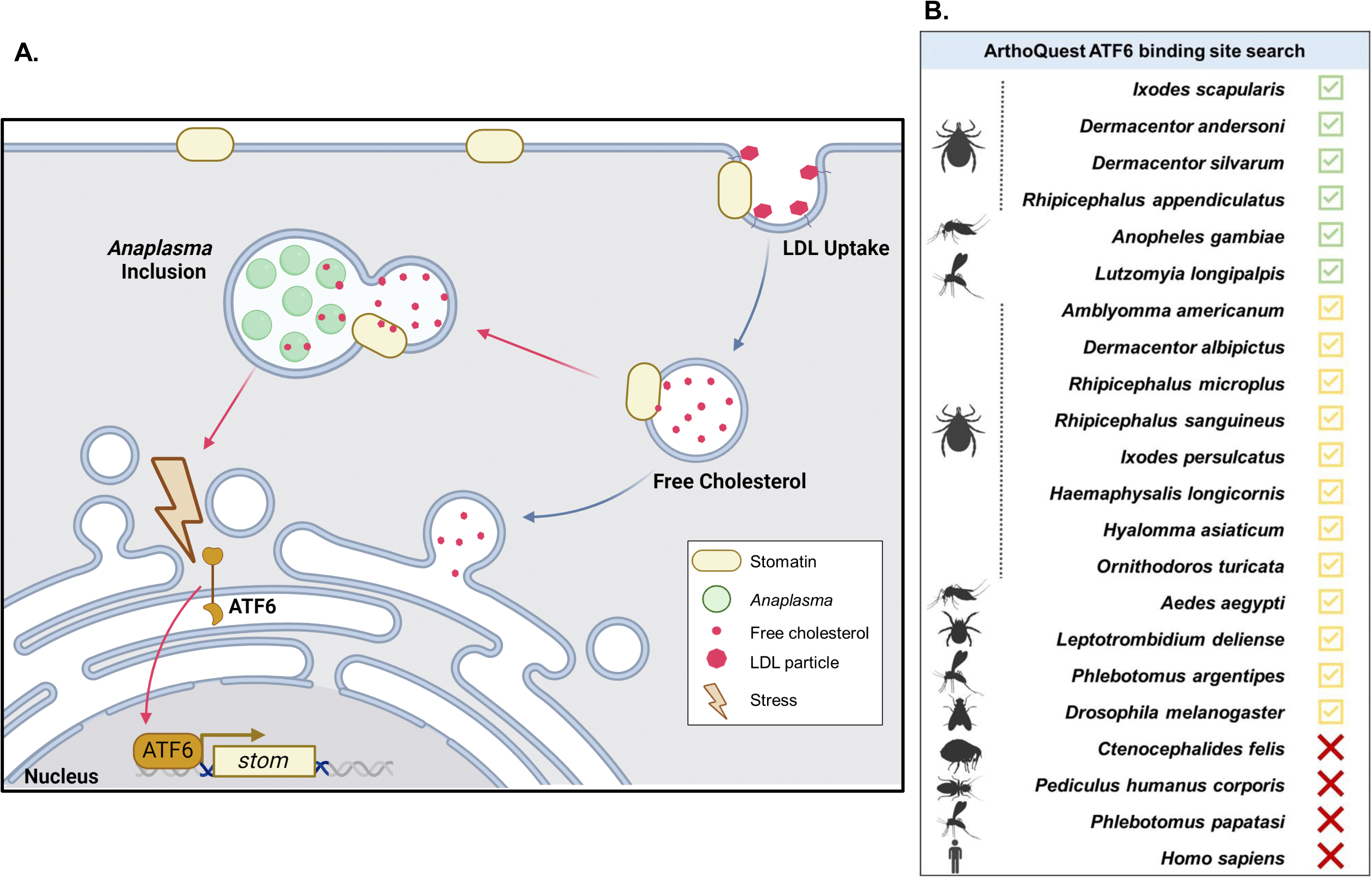
ATF6 supports arthropod infection through Stomatin-regulated cholesterol delivery to the pathogen. (A) Infection activates ATF6 which translocates to the nucleus and upregulates *stomatin* expression. Stomatin regulates cholesterol distribution in the cell and to the *Anaplasma* containing vacuole for growth and survival in ticks. (B) ArthroQuest was used to identify organisms containing ATF6 binding sites in the promoter regions of Stomatin orthologs. Stomatin orthologs were identified using the *Ixodes* protein sequence. Green checkmarks indicate that the top ortholog contains an ATF6 binding site in the promoter region. Yellow checkmarks indicate that a significant ortholog hit contains an ATF6 binding site in the promoter region. Red “X” indicates all BLAST hits lack an ATF6 binding site.

### ArthroQuest: ATF6-regulation of stomatin is unique to arthropod vectors

Programs that predict transcription factor regulatory networks have generally been restricted to model organisms, leaving out most arthropod vectors^41,42^. To address this deficiency, we created a web-based tool termed “ArthroQuest” to serve as a resource for the vector biology community (https://datahub.vetmed.wsu.edu/Shaw/ArthroQuest/). This tool can query 22 pre-loaded arthropod vector genomes for transcription factor binding motifs. Pre-loaded genomes include those from ticks, mosquitoes, lice, sand flies, mites, fleas, *Drosophila*, and humans. Using NCBI genomic FASTA and annotation files, we defined promoter regions for all 22 genomes and created a user-friendly interface with Shiny App. Users can enter a DNA-binding motif of interest and then select a genome to query. Annotated genes found downstream from positive-hit promoter regions can then be downloaded as a table. Using this tool, we queried all pre-loaded genomes for ATF6-binding sites. In contrast to humans, the large majority of arthropods (18 out of 21) had positive hits in the promoter regions of *stomatin* orthologs, including other ticks, mosquitos, and sand flies (Fig 5B, Supplementary Table 4). Altogether, these findings suggest that the ATF6-regulated nature of *stomatin* may be a common feature among blood-feeding arthropods.

## DISCUSSION

Pathogen persistence in vectors can be attributed to both microbe and host-responses, although the underlying mechanisms orchestrating this remain incompletely understood. Recently, the UPR receptors IRE1α and PERK have been connected to arthropod immunity and vector competence of ticks^11,12^. The third pathway orchestrated by ATF6 remains the least understood out of the UPR circuits^13^. In this article, we show that transmissible pathogens *A. phagocytophilum* and *B. burgdorferi* activate ATF6 in *I. scapularis,* which supports pathogen colonization and persistence in the tick. We provide evidence that the ATF6-regulated transcriptional network supports pathogen survival by inducing Stomatin, which facilitates cholesterol delivery to the bacterium. Moreover, queries using our custom, web-based transcription factor binding site tool ArthroQuest suggest that the ATF6-regulated nature of *stomatin* is unique to arthropods. Collectively, our findings provide mechanistic insight into how cellular stress responses influence vector competence of arthropods and further highlight the fundamental differences in molecular networks between mammals and ticks.

To our knowledge, this is the first study to mechanistically investigate how ATF6 influences pathogen dynamics in an arthropod vector. The ATF6 regulatory network controls the expression of genes involved in maintaining protein homeostasis, misfolded protein degradation^16,43^, development, tissue homeostasis, and cytoprotection^14,43^. Our R script querying the *I. scapularis* genome not only predicted known ATF6-regulated genes, but also many others that have not been previously implicated in the ATF6 regulome. This approach led us to the novel finding that ATF6 transcriptionally regulates tick *stomatin*. In addition to *stomatin,* there are several other genes that are unique to the *Ixodes* ATF6 regulome that are not yet characterized (Supplementary Table 2). How these unique targets may be impacting vector competence is an area for future study.

ATF6 has previously been connected to lipid metabolism in mammals^44^. *Ixodes* ticks only feed once per life stage and must adapt to an influx of proteins and lipids when taking a blood meal, which can confer stress to the tick^16^. It is possible that stress-responses are activated during blood feeding and that the ATF6-regulated nature of *stomatin* in ticks is needed for the proper coordination of lipid homeostasis and metabolism. Increased Stomatin expression could stabilize lipid rafts, improve membrane integrity, and/or protect the cell membrane against oxidative stress that is associated with a blood meal. This linkage between ATF6 and lipid homeostasis through Stomatin in arthropods may be something that vector-borne pathogens target and actively manipulate to satisfy their own cholesterol requirements for growth and survival^15^.

Our script in R led to the development of a customizable tool, ArthroQuest, to predict transcription factor binding sites in non-model arthropods (https://datahub.vetmed.wsu.edu/Shaw/ArthroQuest/). Other tools, such as Transfac or TFtarget, exist but are only available for model organisms^41,45^. In addition to humans and *Drosophila,* ArthroQuest currently allows users to query 20 different arthropod vector genomes. This led to the finding that humans do not have an ATF6 binding site in the predicted promoter region of *stomatin*, but the large majority of vectors genomes on ArthroQuest do. This includes several other species of ticks, mosquitos, and sandflies, which suggests that ATF6-mediated regulation of *stomatin* could be a common phenomenon among blood-feeding arthropods. When considering the rapid nutrient influx and temperature shift that hematophagous arthropods endure during blood feeding, a stress-sensing network to cue lipid metabolism would be evolutionarily advantageous, particularly if Stomatin influences cholesterol homeostasis and/or membrane fluidity^33,37,46,47^.

As a lipid-raft associated protein, Stomatin performs several functions within the cell. It has been implicated in the host cell immune response to fungal pathogens by assisting phagolysosome fusion^48^. Despite this association with immunity, Stomatin has also been shown to promote viral infection. For example, Stomatin assists Hepatitis C virus (HCV) replication by supporting assembly of the viral RNA replicase complexes on detergent-resistant membrane structures^49^. Our study is the first to link Stomatin to bacterial infection and replication. Previous reports have found that another SPFH family protein, Flotillin, facilitates cholesterol transport to the *Anaplasma-*containing vacuole in mammalian cells^50,51^. Our findings show that Stomatin-depleted tick cells have increased cholesterol accumulation within the cells, but decreased cholesterol delivery to the bacterium, suggesting that the role of *Ixodes* Stomatin during *Anaplasma* infection could be similar to mammalian Flotillin. This finding highlights that while the cholesterol requirement for *Anaplasma* growth is conserved between mammalian and tick environments, the molecular host targets that facilitate this process are distinct between the two organisms.

While our findings show that both *A. phagocytophilum* and *B. burgdorferi* infection activate ATF6 transcriptional activity, a recent study reported that ATF6 is not activated in RF/6A cells during *Anaplama* infection^52^. The discrepancy in results may be attributable to differences in cell types and/or experimental approaches used to test for ATF6 activation. Wang *et al*. used a transfected RF/6A cell line expressing recombinant, GFP-tagged ATF6 and nuclear accumulation was assessed by fluorescence microscopy after infection^52^. Our assay used a Luciferase reporter plasmid to quantitatively detect transcriptional activity by endogenous, activated ATF6 in infected HEK 293T cells. Another report observed an increase in cleaved ATF6 by immunoblot in *A. phagocytophilum-*infected THP-1 cells, which also suggests activation^53^. It is possible that ATF6 activation may vary between stages of infection. For example, our group previously reported that there was no difference in *atf6* transcript levels between uninfected and *Anaplasma*-infected *Ixodes* nymphs immediately post-repletion^11^. However, in infected larvae that were rested for 7 days post-repletion, we found increased *atf6* expression (Fig 1C-D). The discrepancy in expression at different timepoints post-repletion could indicate that ATF6 is only activated after *Anaplasma* has established infection and expanded in population within the tick. This hypothesis is in line with our findings that ATF6 facilitates cholesterol delivery to the *Anaplasma*-containing vacuole for bacterial growth and survival.

Many vector-borne pathogens are reliant on host cholesterol for membrane structure and growth. For example, *B. burgdorferi, Ehrlichia* spp., and *Anaplasma* spp. incorporate cholesterol into their outer membranes for structural integrity. Other vector-borne parasites and viruses, such as *Plasmodium* spp. and flaviviruses, are also reliant on host lipids for survival^15^. Based on this knowledge and the data presented herein, we hypothesize that ATF6-regulation of Stomatin expression is a mechanism used by a wide variety of blood-feeding arthropods to control lipid homeostasis during times of stress, which is exploited by vector-borne pathogens to effectively colonize arthropods.

## METHODS

### Cell culture

The *I. scapularis* embryonic ISE6 cells (received as gift from Ulrike Munderloh) were cultured at 32°C with 1% CO_2_ in L15C300 media supplemented with 10% heat-inactivated FBS (Sigma, F0926), 10% tryptose phosphate broth (BD, B260300) and 0.1% lipoprotein bovine cholesterol (MP Biomedicals, 219147680).

The human embryonic kidney cell line, HEK 293T cells were cultured in flasks (Corning, 353136) at either 33°C or 37°C in 5% CO_2_ in Dulbecco’s modified Eagle medium (DMEM; Sigma, D6429) supplemented with 10% heat-inactivated FBS (Atlanta Biologicals, S11550) and 1x Glutamax (Gibco, 35050061).

### Bacteria and animal models

*A. phagocytophilum* strain HZ was cultured in HL60 cells (ATCC, CCL-240) in Roswell Park Memorial Institute 1640 (Cytiva SH30027.LS) medium supplemented with 10% heat-inactivated FBS and 1x Glutamax. Cultures were kept between 1 × 10^5^ and 1 × 10^6^ cells mL^−1^ and maintained at 37°C in the presence of 5% CO_2_. Persistently infected ISE6 cells were cultured in L15C300 media supplemented with 25 µM 4-(2-Hydroxyethyl)piperazine-1-ethane-sulfonic acid (Sigma, H4034-100G), 0.25% sodium bicarbonate (Sigma, S-5761), pH 7.5 in unvented flasks (GeneseeSci, 25-207) at 37°C, 5% CO_2_. *A. phagocytophilum* counts were performed as previously described^11,54^. Host cell-free *A. phagocytophilum* was isolated by syringe lysis with a 27-gauge needle.

*B. burgdorferi* B31 (strain MSK5) was grown in modified Barbour-Stoenner-Kelly II (BSK-II) medium supplemented with 6% normal rabbit serum (NRS; Pel-Freez; 31126-5) at 37°C, 5% CO_2_. Dark-field microscopy was used to monitor the density and growth phase of the spirochetes. Plasmid profiles of *B. burgdorferi* cultures were screened by PCR before infection^55^.

*Escherichia coli* cultures were grown overnight at 37°C with shaking between 230 and 250 rpm in lysogeny broth (LB) supplemented with 100 µg µl^−1^ ampicillin.

*Ixodes scapularis* ticks were acquired at the larval stage from either the Biodefense and Emerging Infectious Diseases Research Resources Repository for the National Institute of Allergy and Infectious Disease at the National Institutes of Health (https://www.beiresources.org/) or from Oklahoma State University (Stillwater, OK, USA). Ticks were maintained with 16:8-h light:dark photoperiods and 95-100% relative humidity at 23°C.

Male C57BL/6 mice were obtained from colonies at Washington State University at ages six to ten weeks old. For *A. phagocytophilum* infection experiments, mice were infected intraperitoneally with 1 × 10^7^ host cell-free bacteria in 100 µl of PBS (Intermountain Life Sciences, BSS-PBS) as previously described^11,54^. *A. phagocytophilum* burdens of each mouse were assessed six days post-infection by collecting 25 to 50 µl of blood from the lateral saphenous vein, as previously described^11,12^. *A. phagocytophilum* burdens were quantified by quantitative PCR (*16s* relative to mouse *β-actin*). For *B. burgdorferi* infections, mice were inoculated intradermally with 1 × 10^5^ low-passage spirochetes. Seven days post-infection, blood was collected from the lateral saphenous vein of each mouse and subcultured in BSK-II medium. The presence of spirochetes were confirmed by dark field microscopy^11,12^. All experiments with mice were carried out according to the guidelines and protocols that are approved by the American Association for Accreditation of Laboratory Animal Care (AAALAC) and by the Office of Campus Veterinarian at Washington State University (Animal Welfare Assurance A3485-01). The mice were housed in an AAALAC-accredited facility at Washington State University in Pullman, WA. The Washington State University Biosafety and Animal Care and Use Committees approved all procedures.

### RNAi silencing and pharmacological treatments

The Silencer siRNA Construction Kit (Invitrogen, AM1620) was used to synthesize silencing RNAs (siRNA) and scrambled RNAs (scRNA). For RNAi knockdown experiments, ISE6 cells were seeded at 1 × 10^6^ cells per well in a 24-well tissue culture plate. siRNA or scRNA (3 µg) were transfected into tick cells with 2.5 µl of Lipofectamine 3000 (Invitrogen, L3000008) for 24 hours. Plates were centrifuged at room temperature for 1 hour at 450 x g and then incubated overnight. The following day, cells were infected with *A. phagocytophilum* (MOI 50) for 18 hours. Cells were collected in TRIzol (Invitrogen, 15596026) for RNA isolation. The Direct-zol RNA Microprep Kit (Zymo; R2062) was used to extract RNA. cDNA was synthesized using the Verso cDNA Synthesis Kit (Thermo Fisher Scientific; AB1453B) using 300 to 500 ng total RNA per reaction. Gene silencing, bacterial burden, and gene expression were assessed by quantitative reverse transcription PCR (qRT-PCR) with iTaq universal SYBR Green Supermix (Bio-Rad, 1725125) (Supplementary Table 5). Cycling conditions were as recommended by the manufacturer.

For pharmacological experiments, ISE6 cells were seeded at 1 × 10^6^ cells per well in a 24-well tissue culture plate and treated with 5-50 µM of AA147 (Focus Biomolecules, 10-3973) for 24 hours. RNA isolation, cDNA synthesis, and qRT-PCR were performed as described above. All data are expressed as means ± standard error of the mean (SEM).

### Protein structure predictions and alignments

*Ixodes* ATF6 and *Ixodes* NF-Y were identified using National Center for Biotechnology Information protein Basic Local Alignment Tool (BLAST) and querying the tick genome with human protein sequences (ATF6: AAB64434.1. NY-F: ALX00018.1). All protein alignments were visualized with JalView^56^. Shading indicates physiochemical property conservation between amino acids. AlphaFold was used to model *Ixodes* ATF6 and *Ixodes* Stomatin^20,21^. Alignments with human orthologs were performed in UCSF ChimeraX^22^. *Ixodes* ATF6 binding to the *Ixodes stomatin* promoter was predicted using AlphaFold3 and visualized in ChimeraX^34^. Sequences used for AlphaFold predictions are found in Supplementary Table 1.

### Predicting ATF6 binding sites in the tick genome

All binding site analyses were conducted using R version 4.2.2 and RStudio 2022.07.2.576^57,58^. The *I. scapularis* genome (GCF_01692078) was obtained from the NCBI database in General Feature Format (GFF) and FASTA format^59^. Putative promoter sites were defined as the region 1000 base pairs upstream from each gene. Coding domain sequence (CDS) coordinates from the GFF file were used to identify promoter end sites. Promoter start sites were identified as 1000 nucleotides from the promoter end site on both sense and antisense DNA strands. Using Biostrings^60^ and seqinR^61^ packages, the nucleotide sequence of each of the predicted promoter regions was obtained from the FASTA file. Using the stringR package^62^, ATF6 binding sites within predicted promoter sites were detected using the following motifs: TGACGTG and CCACG with CCAAT. The resulting data included predicted promoter site coordinates and the corresponding gene annotation. To create ArthroQuest, the R script described above was reformatted to create a user-friendly interface with the R package shiny^63^.

To identify orthologs to the putatively ATF6-regulated genes, corresponding protein accession numbers were used to find amino acid sequences from the *I. scapularis* genome using the Rentrez package^64^. Human (GCF_000001405.40) and *Drosophila* (GCF_000001215.4) proteome files were obtained from NCBI as FASTA files. Using the rBLAST package^65^, each protein of interest was queried against human or *Drosophila* proteins. The top 2 hits for each protein of interest were retrieved. All R scripts are available in the GitHub repository (https://github.com/Shaw-Lab/Vosbigian-et-al-2025)

### Gene Enrichment Analysis Visualization

For pathway analysis, ortholog accessions were queried in Gene Ontology and Reactome databases^25–27^. Using the enrichplot package^66^, pathways that were significantly represented (Supplementary Table 3) were plotted. Adjusted *P*-value is indicated by color, Ratios of enriched genes per total annotated genes are indicated by size.

### Plasmid construction

Primers (Supplementary Table 5) were used to amplify the *stomatin* promoter sequence by PCR from ISE6 DNA. The resulting fragment was cloned into a pTE luciferase plasmid (Signosis, LR-2200) using BgIII. *Ixodes nf-y* and the active region of *atf6* (amino acids 1-365) were synthesized by GenScript. *Ixodes atf6* was cloned into pCMV-HA (New MCS) (gift from Christopher A. Walsh; Addgene plasmid number 32530) using EcoRI and EcoRV. *Ixodes nf-y* was cloned into pCMV/hygro-Negative control vector (SinoBiological; CV005) using HindIII and KpnI. All constructs were confirmed by sequencing (Plasmidsaurus).

### HEK 293T cell transfection

1 × 10^6^ HEK 293T cells were seeded into a 6-well plate. The next day, cells were either singly or co-transfected with pCMV-ATF6-HA and pCMV-NFY-FLAG using 10 µl Lipofectamine 3000, 10 µl P reagent (Fisher Scientific; L30000015), in Opti-MEM I reduced-serum medium (Gibco; 31985062). After 5 hours, the medium was replaced with complete DMEM and cells were incubated at 33°C, 5% CO_2_ for 24 hours. Cells were lysed and collected in RIPA (radioimmunoprecipitation assay; Fisher Scientific; PI89901) supplemented with 1 x protease and phosphatase inhibitors (ThermoScientific; 78440)^11,12^.

### Polyacrylamide gel electrophoresis and Western blotting

Protein concentrations were determined by BCA assay (Bicinchoninic acid assay; Pierce; 23225). For each sample, 20 µg of protein was separated using a 4-15% MP TGX precast cassette (Bio-Rad; 4568084) at 100 V for approximately 1 hour and 30 minutes before being transferred to a PVDF (polyvinylidene difluoride) membrane. Membranes were blocked with 5% milk in 1 x PBS-T (phosphate-buffered saline containing 0.1% Tween 20) for approximately 1 hour at room temperature. All primary antibodies were diluted in 5% milk, PBS-T and incubated with the blot overnight at 4°C. The primary antibodies used were anti-HA anti-mouse (Invitrogen 26153; 1:1000), anti-FLAG-HRP diluted (Sigma A8512; 1:2000), and anti-Actin (Sigma A2105; 1:1000). Membranes were then washed 3 times with 0.5% milk, PBS-T before adding a secondary antibody. Secondary antibodies included rabbit anti-mouse (Star13B; 1:2000 dilution) and donkey anti-rabbit (Sigma; A16023; 1:5000). Secondary antibodies were incubated with membranes for 2 hours at room temperature. Membranes were then washed and imaged using an ECL western blotting substrate (Enhanced chemiluminescence; Fisher Scientific; P13216).

### Luciferase Reporter Assays

For the ATF6 activation assay, 1 × 10^4^ HEK 293T cells were seeded into white-walled, clear-bottom 96-well plates (Greiner Bio-One, 655098). The following day, cells were transfected with 0.05 µg of the ATF6 luciferase reporter plasmid using 0.5 µl of Lipofectamine 3000 in Opti-MEM I reduced-serum medium (Gibco, 31985062). Transfections were allowed to proceed overnight. The following day, cells were infected for 18 hours with *A. phagocytophilum* (MOI 50) or *B. burgdorferi* (MOI 200) or left uninfected. Luminescence was measured the following day by adding 5 mg mL^−1^ of D-Luciferin potassium salt (Promega, E1500) to each well and quantifying with a plate reader. Data is represented as relative luciferase units (RLU) normalized to non-infected controls ± SEM.

To assess *stomatin* activation by ATF6, cells were triple-transfected with the reporter plasmid containing Luciferase under control of the *stomatin* reporter and plasmids expressing *Ixodes* ATF6 and NF-Y (0.05 μg of each). Luminescence was measured as described above. Data is normalized to the control containing only the *stomatin* reporter plasmid without ATF6 and NF-Y expression plasmids.

### Gene expression analysis of whole ticks

Gene expression profiling was performed on ticks at both larval and nymph stages. Replete larvae were collected after being fed on either clean mice, an *A. phagocytophilum*-infected mouse, or *B. burgdorferi-*infected mouse. *A. phagocytophilum-*infected larvae were collected and maintained in an incubator for either 7 days or were allowed to molt to nymphs. *B. burgdorferi-*infected larvae were collected and either maintained for 14 days post-repletion or were allowed to molt to nymphs. When isolating RNA, individual ticks were flash frozen in liquid nitrogen and mechanically pulverized before adding TRIzol. RNA was isolated and cDNA was synthesized as described above. Primers listed in Supplementary Table 5 were used to measure gene expression by qRT-PCR as described above. All samples were normalized to uninfected controls. Data is expressed as means ± SEM.

### RNAi silencing and analysis of whole ticks

*I. scapularis* larvae were silenced with RNAi as previously described^11,12^. Around 150 larvae were placed in a 1.5 mL tube with 40 µl of siRNA or scrambled controls and incubated overnight at 15°C. Larvae were then dried and allowed to recover overnight before being placed on infected mice. Replete larvae were collected over a period of 3-5 days and frozen. To assess feeding efficiency, larvae were weighed in groups of three. RNA was isolated from individual ticks as described above. To generate absolute numbers of the target sequences, qRT-PCR was performed with a standard curve. Standard curves were generated with a plasmid containing either *A. phagocytophilum 16s, B. burgdorferi flab, Ixodes β-actin, or Ixodes stomatin* (Supplementary Table 5).

### Cholesterol Accumulation Assays

To quantify cholesterol in tick cells, the Amplex Red Cholesterol Assay Kit (Invitrogen, A12216) was used. 6.5 × 10^7^ of ISE6 cells were seeded in T-25 tissue culture flasks (Greiner Bio-one 07-000-226). The next day, the monolayer was washed three times with PBS and resuspended in 1 mL of PBS. An aliquot was set aside for RNA isolation and *β-actin* measurement. The rest of this sample was used for cholesterol quantification. 50 µl of each sample was added to a 96 well clear-bottom, black-sided plate (Thermo Scientific, 12-566-70). Total cholesterol was quantified via the Amplex Red Cholesterol Assay Kit (Invitrogen, A12216) according to the manufacturer’s instructions and absorbance was measured at 590 nm. A standard curve was used to calculate total cholesterol concentration. Data is normalized to absolute copies of *β-actin*.

To quantify cholesterol in *A. phagocytophilum,* 4 × 10^6^ cells of persistently infected ISE6 cells that were silenced for *stomatin* (12 µg of siRNA or scRNA with 10 µl Lipofectamine 3000) were seeded into 6-well plates (STARSTEDT, 83.1839). 5 days post-transfection cells were collected in PBS. Host cell-free *A. phagocytophilum* were isolated by sonication. Briefly, cells were centrifuged at 2300 x g for 10 minutes at 4°C and resuspended in 500 µl of PBS. Samples were sonicated four times for 15 seconds at an amplitude of 30V and lysates were centrifuged at 710 x g for 5 minutes to remove host cell debris. The supernatant was collected and an additional spin at 2300 x g for 10 minutes was performed to collect *A. phagocytophilum*. Samples were resuspended in 200 µl of PBS. 100 µl was used to quantify cholesterol as described above. The remaining sample was used to quantify bacteria by isolating RNA, synthesizing cDNA, and quantifying *phagocytophilum 16s* by qRT-PCR. Total cholesterol was normalized to *A. phagocytophilum 16s.* One well was collected in TRIzol and analyzed by qRT-PCR to assess silencing efficiency.

### Statistical analysis

*In vivo* experiments used at least 10-20 ticks and *in vitro* experiments had at least three to five replicates. Data was analyzed with a non-parametric Mann-Whitney test or an unpaired Student’s t-test respectively and expressed as means ± SEM. GraphPad Prism was used for calculations and creating graphs. Statistical significance was determined by a *P* value of <0.05.

## ACKNOWLEDGEMENTS

We are grateful to Ulrike Munderloh (University of Minnesota) for providing ISE6 tick cell lines, Jon Skare (Texas A&M Health Science Center) for providing *B. burgdorferi* B31 (MSK5), Skandha Nadarajah, Ryan Driskell, and the Veterinary Information Systems team (Washington State University) for their assistance with making ArthroQuest publicly available, Ryan Vosbigian (University of Idaho) for his guidance on R programming, Biodefense and Emerging Infectious Diseases Resources and Oklahoma State University for *Ixodes scapularis* ticks. The Addgene plasmid 32530 was a gift from Christopher A. Walsh. Schematics in Fig 1 and 7 were created with BioRender.

## Author Contributions

K.A.V and D.K.S. designed the study. K.A.V., S.J.W., K.L.R., E.A.F, E.R-Z., and D.K.S. performed experiments. K.A.V. and E.A.S. developed ArthroQuest. K.A.V. and D.K.S. analyzed data. All authors provided intellectual input into the study. K.A.V., S.J.W., and D.K.S. wrote the manuscript. All authors contributed to editing.

