## Supplementary Figure 1 for "ATF6 enables pathogen infection in ticks by inducing *stomatin* and altering cholesterol dynamics"

**<u>Supplemental Figure 1.</u> *I. scapularis* ATF6 sequence alignment and structural prediction with AlphaFold.**

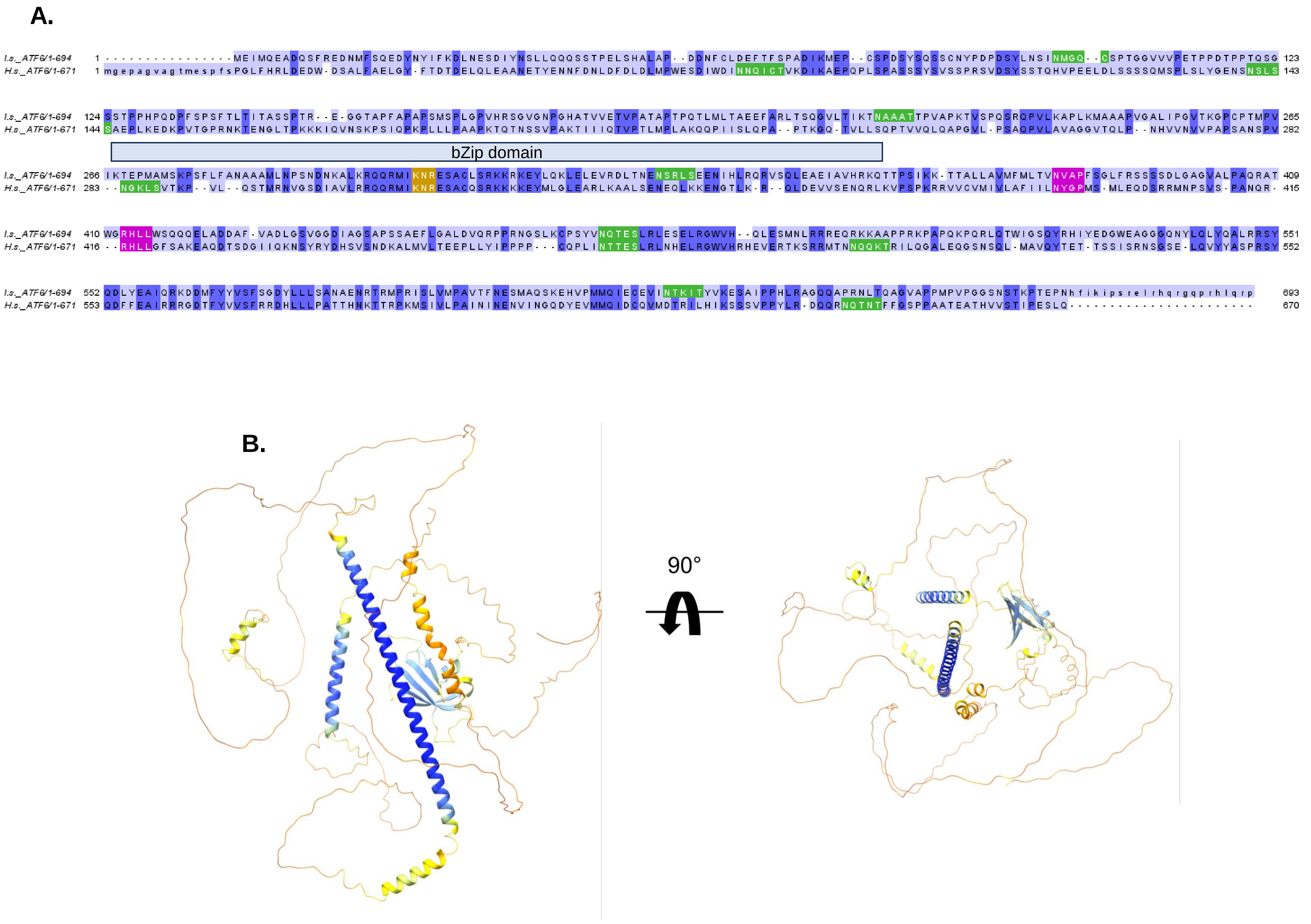
