## Supplementary figures and images for "ATF6 enables pathogen infection in ticks by inducing *stomatin* and altering cholesterol dynamics"

### Supplementary Figure 2

**<u>Supplementary Figure 2.</u> Predicted ATF6 regulatory network across organisms.**

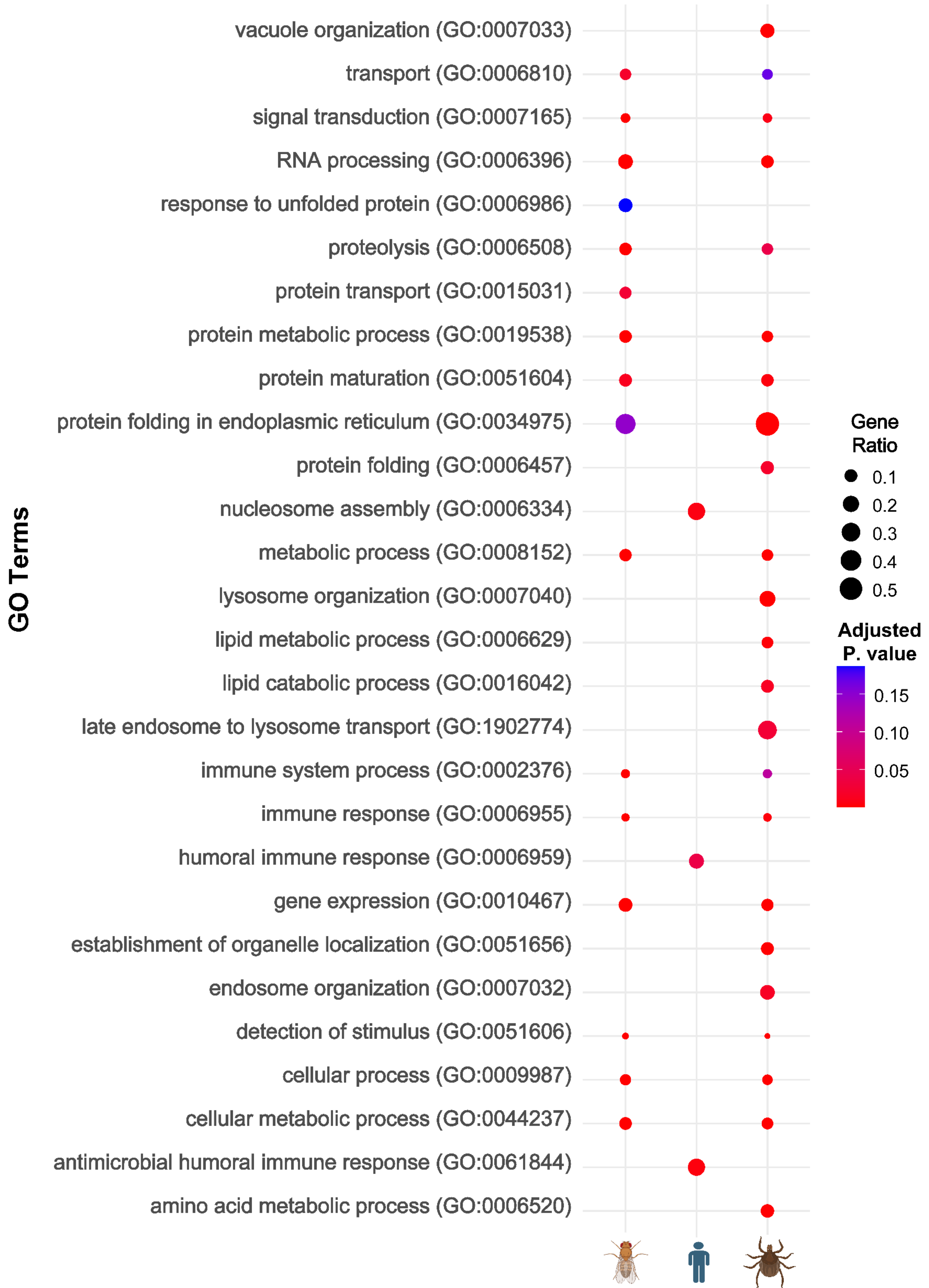

### Supplementary Figure 4

**Supplementary Figure 4.** ATF6 schematic and recombinant protein expression with NF-Y

A.

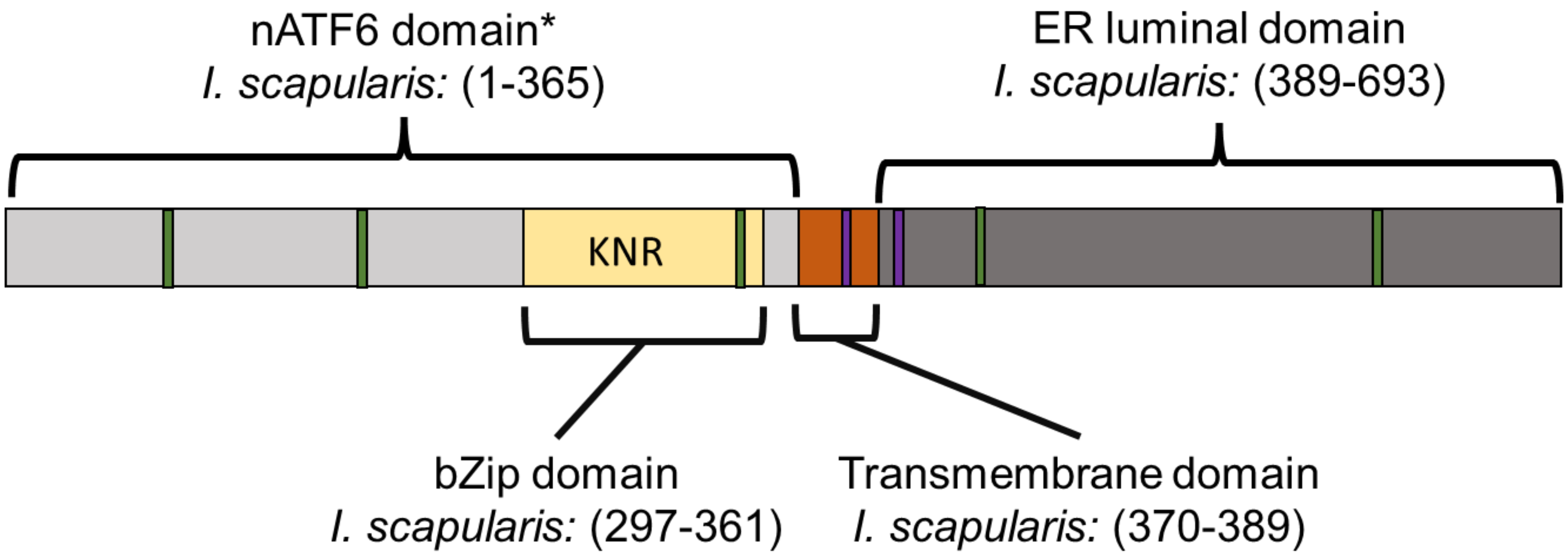


B.

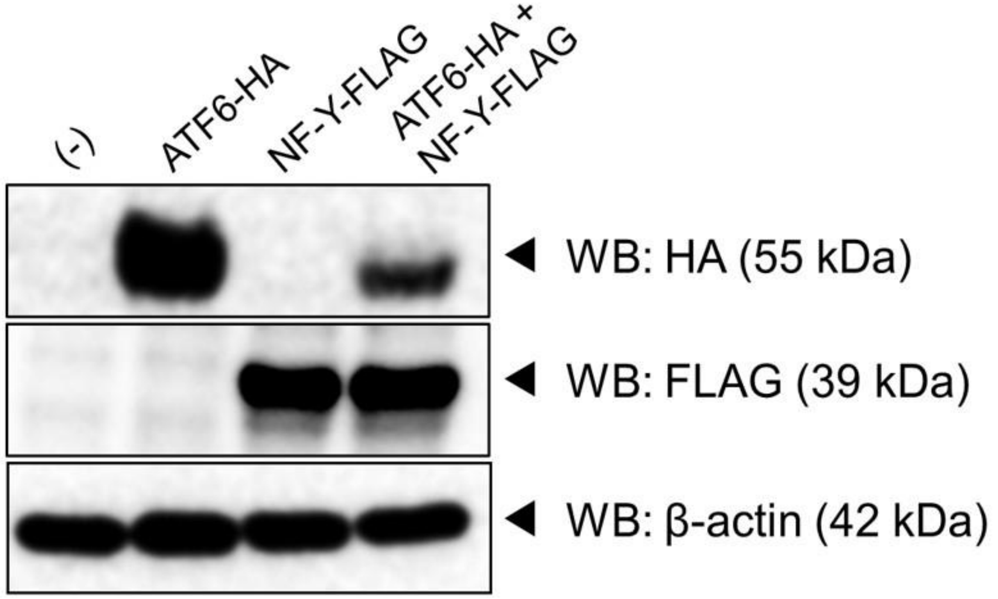


C.

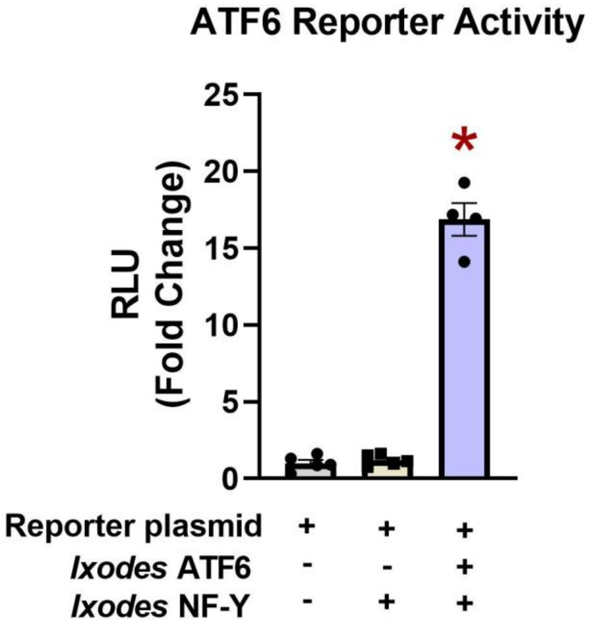
