## Supplementary Figure 3 for "ATF6 enables pathogen infection in ticks by inducing *stomatin* and altering cholesterol dynamics"

**<u>Supplementary Figure 3.</u> Structural interaction of ATF6 and cell viability post AA147 treatment.**

A.

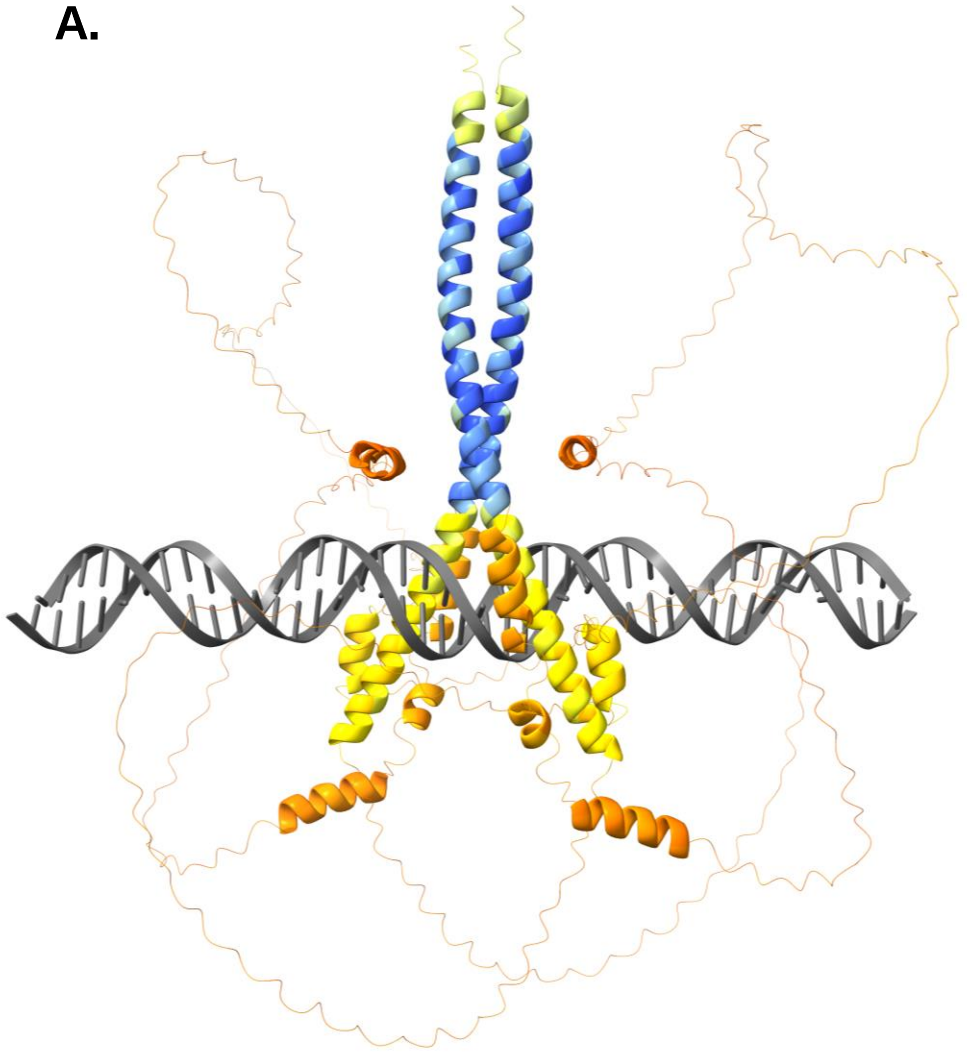

B.

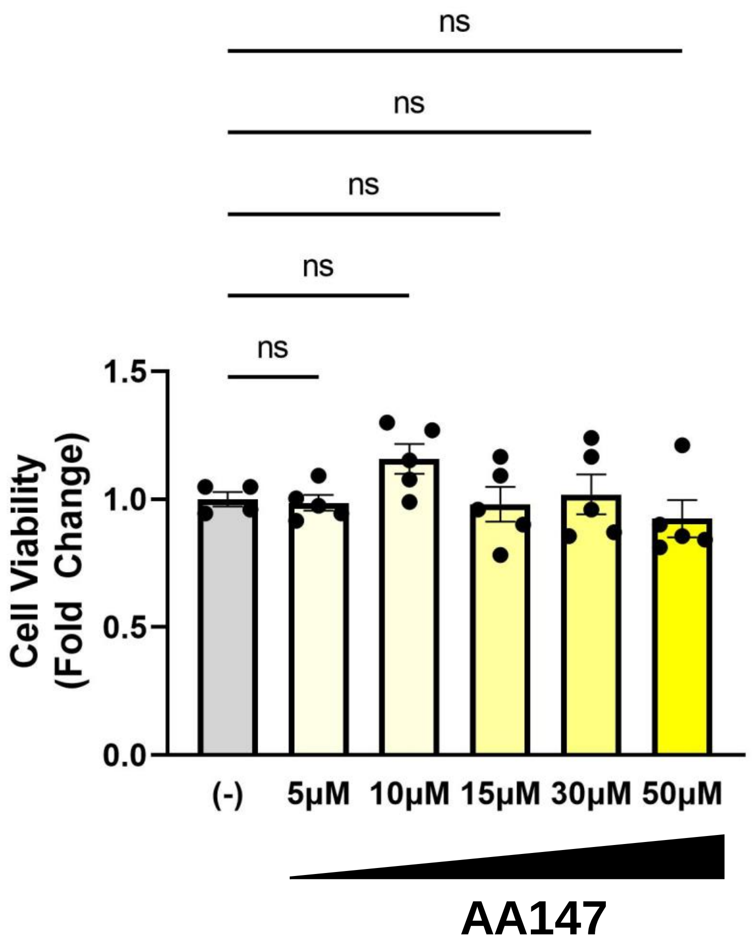
