## Supplementary Figure 5 for "ATF6 enables pathogen infection in ticks by inducing *stomatin* and altering cholesterol dynamics"

### Supplemental Figure 5. *I. scapularis* Stomatin sequence, structural, and domain analysis

**A.**

```
I.s._STOM/1-262        1 MPSPVPD-HPFGVVLKVISLFLIVITLPFSLLLCLVVVQEFERAVIFRLGRLQPGGAAGPGLFFII  65
Human_STOM/20-282     20 KDSPSKGLGPCGWILVAFSFLFTVITFPISIWMCIKIIKEYERAIIFRLGRILQGGAKGPGLFFIL  85

I.s._STOM/1-262       66 PCIDEYRVVDLRTVVFNVCPQEILSKDSVTVAVDAVVYYRVFNPVAATVNIKDHARSTILLAATIL 131
Human_STOM/20-282     86 PCTDSFIKVDMRTISFDIPPQEILTKDSVTISVDGVVYYRVQNATLAVANITNADSATRLLAQTTL 151

I.s._STOM/1-262      132 RNVLGTKMLSDVLSQRESISRTMQTLLDVATDPWGVKVERVELTDVQLPAQMQRAMAAEAEAVREG 197
Human_STOM/20-282    152 RNVLGTKNLSQILSDREEIAHNMQSTLDDATDAWGIKVERVEIKDVKLPVQLQRAMAAEAEASREA 217

I.s._STOM/1-262      198 RAKVVAAEGEQRAAVALRNAANVIAQSPAALQLRYLQTLGTISAEKNSTIVFPLPLELLQTLLFR  262
Human_STOM/20-282    218 RAKVIAAEGEMNASRALKEASMVITESPAALQLRYLQTLTTIAAEKNSTIVFPLPIDMLQGIIGA  282
```

**B.**

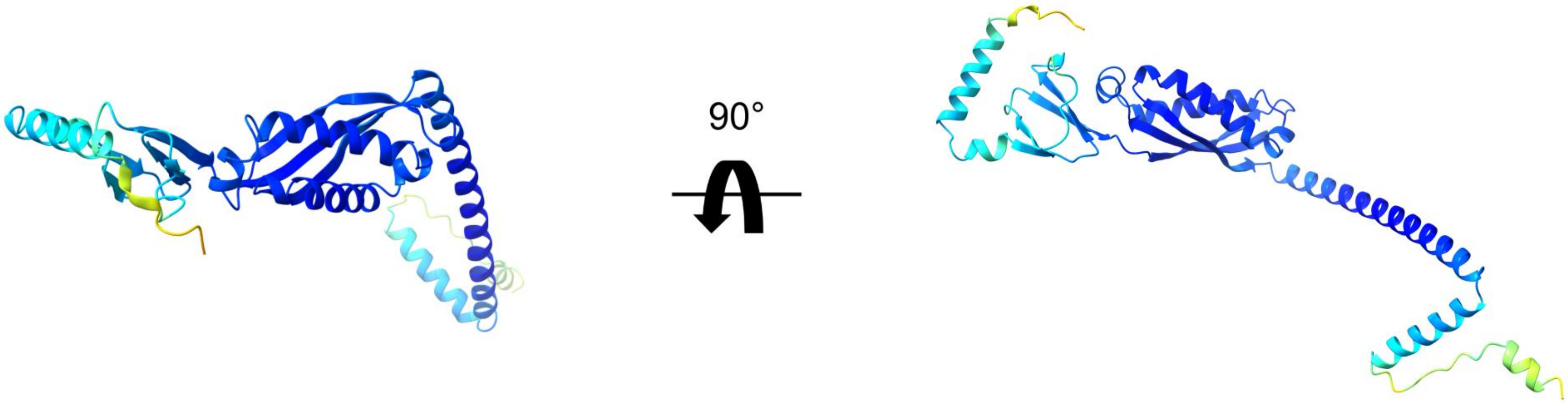


**C.**

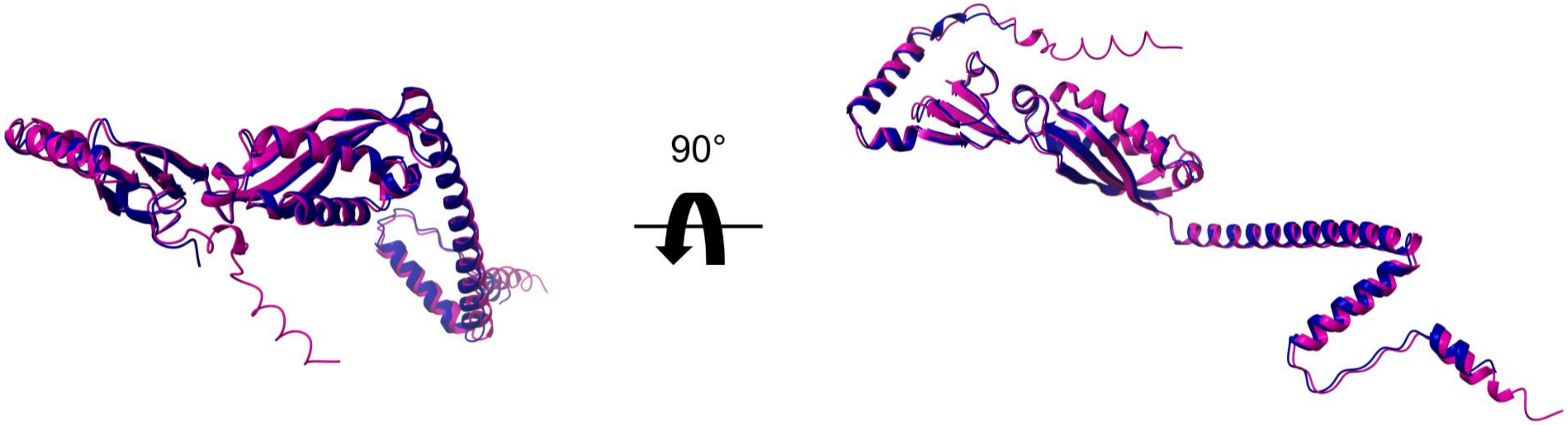

**Supplemental Figure 5.** ***I. scapularis*** **Stomatin sequence, structural, and domain analysis**

**D.** **Cholesterol binding motifs on *Ixodes* Stomatin**

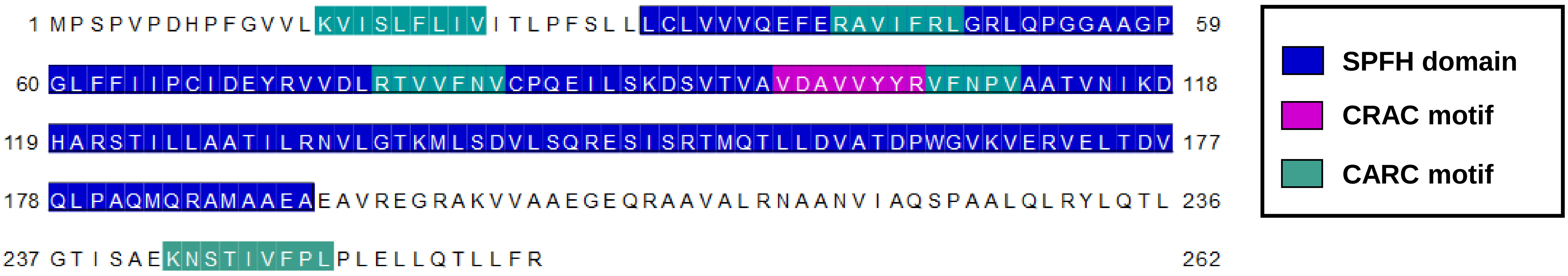


**E.**

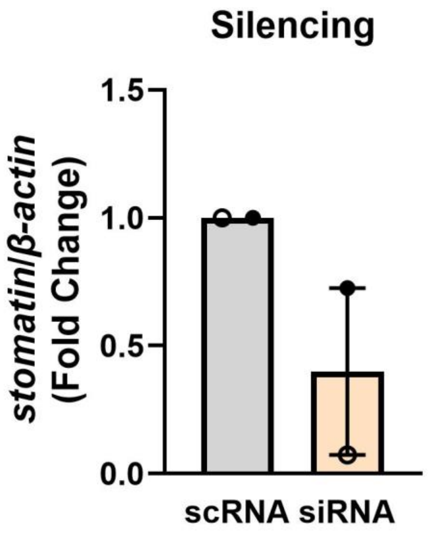
