## Supplementary Table 5 for "ATF6 enables pathogen infection in ticks by inducing *stomatin* and altering cholesterol dynamics"

**Supplemental Table 5.** Oligonucleotide primers used in this study

| **Name** | **Target gene** | **Primer Sequences** |
| --- | --- | --- |
| Mus musculus β-Actin (qRT-PCR) | XM_030254057.1 | F 5'-ACGCAGAGGGAAATCGTGCGTGAC-3'  R 5'-ACGCGGGAGGAAGAGGATGCGGCAGTG-3' |
| Anaplasma phagocytophilum 16S (qRT-PCR) | NC_007797 | F 5'-CCCTAAGGCCTTCCTCACTC-3'  R 5'-CAGCCACACTGGAACTGAGA-3' |
| Anaplasma phagocytophilum 16S_full | NC_007797 | F 5'- TCCTGGCTCAGAACGAACG-3'  R 5'- GTCACTGACCCAACCTTAAATGG-3' |
| Borrelia burgdorferi FlaB (qRT-PCR) | MN954474.1 | F 5'-TTGCTGATCAAGCTCAATATAACCA-3'  R 5'-TTGAGACCCTGAAAGTGATGC-3' |
| Ixodes scapularis Actin (qRT-PCR) | XM_029977298.1 | F 5'-GCCGGGACCTTACAGACTATC-3'  R 5'-CACGGACAATTTCACGCTCG-3' |
| Ixodes scapularis ATF6 (qRT-PCR) | XM_029979488.3 | F 5’-TGCTGACAGCGGAAGAGTTT-3’  R 5’-TTAAGGGGAGCCTTGAGCAC-3’ |
| Ixodes scapularis Stomatin (qRT-PCR) | XM_002412522.3 | F 5’-CCCTCTTCCTCATCGTCATCA-3’  R 5’-TCGAATTCTTGCACGACGAC-3’ |
| Borrelia burgdorferi cp9 (PCR) | BBC10 | F 5'-GAACTATTTATAATAAAAAGGAGAGC-3'  R 5'-ATCTTCTTCAAGATATTTTATTATAC-3' |
| Borrelia burgdorferi cp26 (PCR) | BBB19 | F 5'-AATAATTCAGGGAAAGATGGG-3'  R 5'-AGGTTTTTTTGGACTTTCTGCC-3' |
| Borrelia burgdorferi Ip17 (PCR) | BBD10 | F 5'-CAAACTTATCAAATAGCTTATC-3'  R 5'-ACTGCCACCAAGTAATTTAAC-3' |
| Borrelia burgdorferi Ip25 (PCR) | BBE16 | F 5'-ATGGGTAAAATATTATTTTTTGGG-3'  R 5'-AAGATTGTATTTTGGCAAAAAATTTTC-3' |
| Borrelia burgdorferi Ip28-1 (PCR) | BBF20 | F 5'-ATGAACAAAAAATTTTCTATTTC-3'  R 5'-GTTGCTTTTGCAATATGAATAGG-3' |
| Borrelia burgdorferi Ip28-2 (PCR) | BBG02 | F 5'-TCCCTAGTTCTAGTATCTACTAGACCG-3'  R 5'-TTTTTTTTGTATGCCAATTGTATAATG-3' |
| Borrelia burgdorferi Ip28-3 (PCR) | BBH06 | F 5'-GATGTTAGTAGATTAAATCAG-3'  R 5'-TAATAAAGTTTGCTTAATAGC-3' |
| Borrelia burgdorferi Ip28-4 (PCR) | BBI16 | F 5'-CAGGCCGGATTTTAATATCGA-3'  R 5'-GTTTATATTTTGACACTATAAG-3' |
| Borrelia burgdorferi Ip36 (PCR) | BBK19 | F 5'-AAGTTTATGTTTATTATTGC-3'  R 5'-ATTGTTAGGTTTTTCTTTTCC-3' |
| Borrelia burgdorferi Ip38 (PCR) | BBJ34 | F 5'-AAATTCTATGGAAGTGATG-3'  R 5'-TTTCTATTTATTTTTAGGC-3 |
| Borrelia burgdorferi Ip54 (PCR) | BBA16 | F 5'-GCACAAAAAGGTGCTGAG-3'  R 5'-TTTTAAAGCGTTTTTAAGC-3' |
| Borrelia burgdorferi Ip56 (PCR) | BBQ56 | F 5'-AAGATTGATGCAACTGGTAAAG-3'  R 5'-CTGACTGTAACTGATGTATCC-3' |
| Ixodes scapularis ATF6 siRNA_69 | XM_029979488.3 | F 5’-AAGCTGACCATTAAGACGAATCCTGTCTC-3’  R 5’-AAATTCGTCTTAATGGTCAGCCCTGTCTC-3’ |
| Ixodes scapularis ATF6 scRNA | N/A | F 5’-AAGTATTACCCAACGAGAGTACCTGTCTC-3’  R 5’-AAGCTATTGGCTTGCTTAACACCTGTCTC-3’ |
| Ixodes scapularis Stom siRNA_182 | XM_002412522.3 | F 5’-AAGTCCGGGTCTATTCTTCATCCTGTCTC-3’  R 5’-AAATGAAGAATAGACCCGGACCCTGTCTC-3’ |
| Ixodes scapularis Stom scRNA | N/A | F 5’-AAGCTATTGGCCTTAGTCTCTCCTGTCTC-3’  R 5’-AAAGAGACTAAGGCCAATAGCCCTGTCTC-3’ |
| pCMV-ATF6-HA | XM_029979488.3 | F 5’-AAAAGAATTCATGGAAATTATGCAGGAGGCTGATC-3’  R 5’- AATTGATATCTCATTTTTTAATGCTGGGGGTGGTC-3’ |
| pCMV-NFY-FLAG | XM_029991788.4 | F 5’-AACCAAGCTTATGCAGATGGAGCAGTACCAGGT-3’  R 5’-ACTTGGTACCGTTGGAGTCGCCCCCAGC-3’ |
| Pte-StomPromoter-Luciferase | Predicted promoter region found in Table S2 | F 5’-ACGCAGATCTTAACGTGCATCACGCAGT-3’  R 5’-ACGCAGATCTTCCCTTCGTTTCCTCCTCTTTG-3’ |
